# Soil Microbial Diversity of the Imperial Palace Outer Gardens, Tokyo

**DOI:** 10.64898/2026.03.30.715141

**Authors:** Honami Ando, Runa Furuya, Kohei Ito

## Abstract

The Imperial Palace in Tokyo serves as a significant reservoir of biodiversity within the urban landscape; however, its soil microbial communities remain uncharacterized despite decades of macro-biological surveys. This study presents the first dataset profiling the soil microbiome of the Imperial Palace Outer Gardens, utilizing both 16S rRNA amplicon and shotgun metagenomic sequencing to fill this knowledge gap. We collected bulk soil samples from four distinct sites, including pond sediments and soils beneath ginkgo and pine trees, to capture a range of environmental conditions within this conserved greenspace. Both 16S rRNA amplicon sequencing and shotgun metagenomic sequencing revealed that Pseudomonadota and Actinomycetota were the predominant phyla across all samples. Notably, sites with monoculture vegetation, such as those beneath pine trees, exhibited lower microbial diversity than other locations. Functional annotation identified core metabolic pathways and detected specific antimicrobial resistance and virulence factor genes in selected samples. These datasets provide a critical baseline for future research into urban ecosystem dynamics, soil health, and the intersection of environmental conservation and public health.

## Introduction

The Imperial Palace in Tokyo encompasses an area of 2.3 km^2^, surrounded by approximately 6,897 meters of moats. Much of the area consists of forests, grasslands, moats, and ponds, providing an important natural environment within the urban center. This rich natural landscape has been conserved by the Imperial Household Agency and the Ministry of the Environment, and continuous biodiversity surveys have been conducted since 1996. During Phase I of the survey (1996–2000), 1,366 plant species and 3,638 animal species were recorded. In the subsequent Phase II survey (2009–2014), an additional 250 plant and 649 animal species were identified, including numerous newly discovered, threatened species, as well as species previously thought to be extinct within the Tokyo metropolitan area [1–5]. Many species with later migration periods, such as birds, have also been observed, suggesting that the Imperial Palace’s ecosystem interacts with the surrounding environment [6].

Although the presence of diverse flora and fauna in the Imperial Palace has been well documented, no studies have reported on the microorganisms that form the foundational layer of these ecosystems. Consequently, the composition and structure of the microbial ecosystems in this environment remain unknown.

Microbial ecosystems and their diversity are not only the foundation of ecosystems but also play a vital role in urban public health [7,8]. Exposure to diverse microorganisms is known to contribute to the maturation of the human immune system and to the suppression of infectious disease outbreaks. Multiple previous studies have suggested that childhood exposure to diverse environmental microbiota supports immune development and reduces the risk of allergies and respiratory diseases later in life [9–13]. In fact, it has been reported that microorganisms are transmitted to humans through publicly accessible green spaces such as the Imperial Palace [14,15]. In this study, we collected soil samples from the Imperial Palace and conducted 16S rRNA amplicon sequencing and shotgun metagenomic sequencing to characterize the soil microbial community structure and its functional diversity. Elucidating the microbial diversity within this conserved yet urban environment may provide valuable insights for developing urban design policies that address sustainable and public health challenges.

## Material and methods

### Sampling

Bulk soil samples were collected on 21 October 2025 from the Outer Gardens of the Imperial Palace and Kitanomaru Park, Tokyo, Japan. Samples were obtained from four distinct locations: (1) sediment from a pond in Kitanomaru Park, where the surrounding area was planted with tree species such as Chinese elm (*Ulmus parvifolia*), camphor tree (*Cinnamomum camphora*), Japanese castanopsis (*Castanopsis sieboldii*), Japanese zelkova (*Zelkova serrata*), and Japanese bayberry (*Morella rubra*). Additionally, an accumulation of leaf litter was observed on the pond bed (2) soil beneath a ginkgo tree in Kitanomaru Park. The surrounding area was planted with tree species such as ginkgo (*Ginkgo biloba*), false daphne (*Daphniphyllum macropodum*), and camphor tree (*Cinnamomum camphora*). The ground was covered with various types of undergrowth, where an accumulation of ginkgo seeds and leaf litter was observed, (3) soil beneath a pine tree near the Sakashita Gate in the Outer Gardens. The surrounding area was planted with Japanese black pine (*Pinus thunbergii*), and the ground appeared to be covered by a monoculture of lawn grass. Unlike other sites, there was no significant accumulation of leaf litter, and the soil was notably firmer than in the other samples, and (4) soil from the moat near the Sakuradamon Gate in the Outer Gardens. Although the surrounding trees consisted solely of Japanese black pine (*Pinus thunbergii*), the ground was covered with diverse undergrowth, and an accumulation of leaf litter was present.

Pictures of each site where samples were collected are shown in **Figure 1**. At each site, two subsamples were collected from a depth of 10 cm below the surface. The sterile cotton-tipped swab (ESwab™; Copan Diagnostics, Brescia, Italy) was used for soil collection. A hole approximately 10 cm deep was dug in the ground, and the swab was inserted into the hole to collect the soil adhering to it, then stored in a tube containing Liquid Amies Medium solution. Sampling procedures were designed to minimize contamination and avoid disturbance of the native microbial communities.

**Figure 1.**
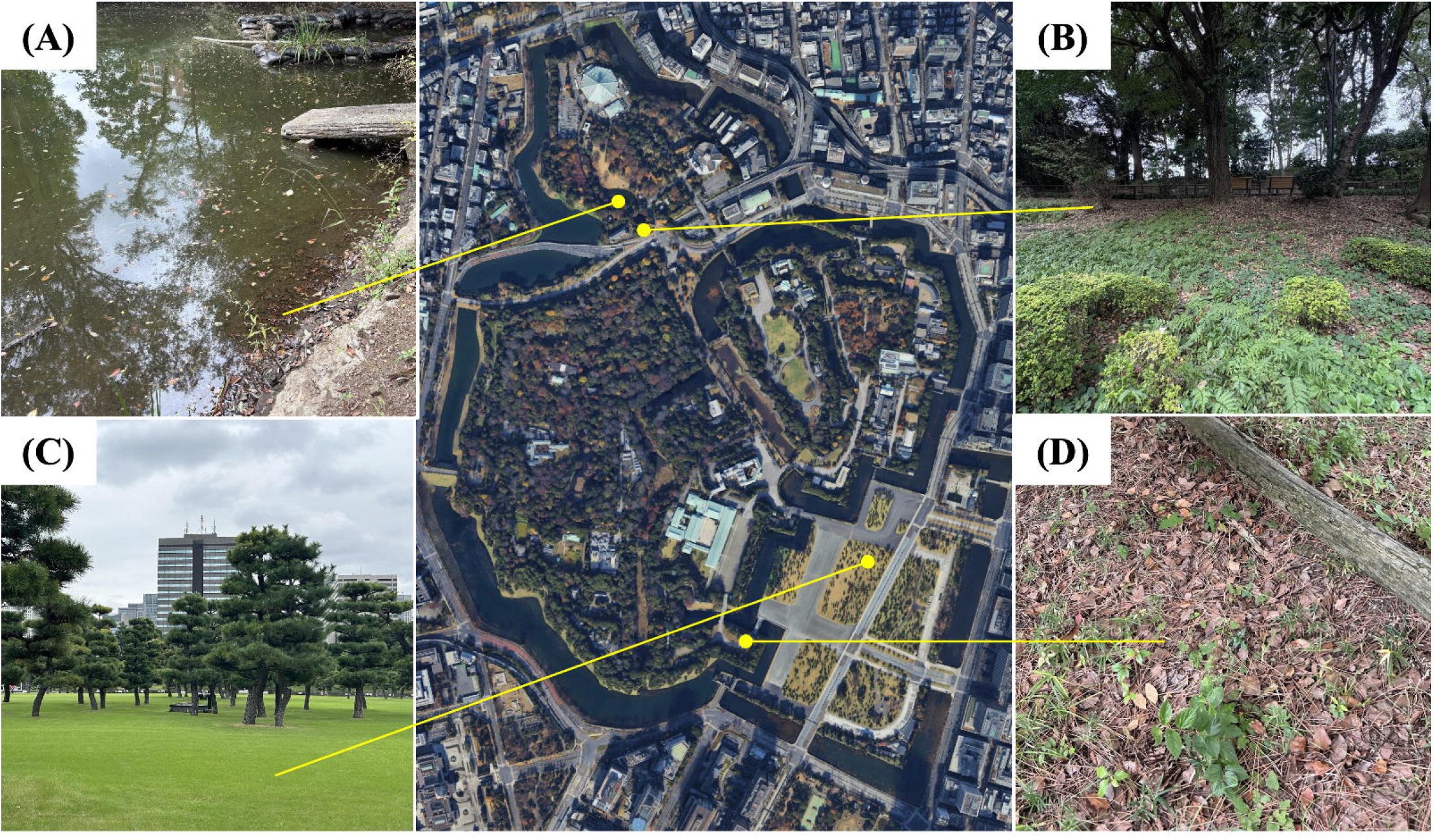
Pictures of the sampling locations. (A) KOKYO_01 and KOKYO_01_2; sediment from a pond in Kitanomaru Park, (B) KOKYO_02 and KOKYO_02_2; soil beneath a ginkgo tree in Kitanomaru Park, (C) KOKYO_03 and KOKYO_03_2; soil beneath a pine tree near the Sakashita Gate in the Outer Gardens, (D) KOKYO_04 and KOKYO_04_2; soil from the moat near the Sakuradamon Gate in the Outer Gardens

### Environmental profile

The Imperial Palace and the Outer Gardens encompass 2.3 km^2^ sites, surrounded by a moat measuring 6,897 meters in length. The Outer Gardens cover an area of 1.15 km^2^ and consist of four areas: the Outer Gardens district, centered around the Imperial Palace Plaza; the Kitanomaru district located on the north side of the Imperial Palace; and the Outer Perimeter district, which comprises twelve moats encircling the palace. The average air temperature in Chiyoda Ward, Tokyo, where the Imperial Palace is located, is approximately 18°C in October, with a total monthly precipitation of 234.8 mm.

### Geographic range

1-1 Kokyogaien, Chiyoda-ku, Tokyo, Japan (Latitude/Longitude: E139°45’31.47”, N35°40’28.67”).

### Sample processing

The samples were promptly sealed, immediately frozen and stored at −30°C until DNA extraction. DNA was extracted from all nine samples using the DNeasy® PowerSoil® Pro Kit (QIAGEN, Germany) according to the manufacturer’s protocol. The extracted DNA was quantified using Qubit™ (Thermo Fisher Scientific, Massachusetts, USA).

### 16S rRNA amplicon sequencing

The first PCR cycle was conducted using ExTaq HS DNA Polymerase, where specific primers 341F (5’-CCTACGGGNGGCWGCAG-3’) and 805R (5’-GACTACHVGGGTATCTAATCC-3’) of targeting the V3–V4 region of the 16S rRNA were amplified [16]. Two microliters of the first PCR product, purified using AMPure XP beads, served as a template for library preparation and the second PCR product were again purified using AMPure XP beads. 16S rRNA gene amplicon sequencing was performed using 460 bp paired-end sequencing on the Illumina MiSeq platform (Illumina Inc., San Diego, CA, USA) at FASMAC Co., Ltd. (Kanagawa, Japan).

### Shotgun metagenome sequencing

Sequencing libraries were prepared for the shotgun metagenomic sequencing, and quality control (QC) was performed at each step to ensure data reliability. Quantified libraries were prepared using KAPA EvoPlus V2 Kit (42 ng input). The libraries were then pooled and sequenced using the DNBSEQ platform (MGI Tech Inc.,Shenzhen,China) according to the effective library concentration and the required data amount, producing a 150-bp paired-end indexed recipe at FASMAC Co., Ltd. (Kanagawa, Japan).

### Data processing

#### Microbiome analysis of 16S rRNA amplicon sequences

Microbiome analysis was performed based on previous studies [17,18].Briefly, raw FASTQ data files were imported into the QIIME2 platform (2024.10) as qza files [19] for quality control and denoising using DADA2 [20]. The sequences were organised into amplicon sequence variants (ASVs), which were then classified against the SILVA SSU 138 database using the QIIME feature-classifier classification scikit-learn package [21,22]. The subsequent analysis excluded ASVs classified as mitochondria, chloroplast, or unassigned. A subsample of 5,000 reads was taken to reduce bias caused by differences in read depth between samples. Shannon entropy, the number of Faith’s phylogenetic diversity were calculated, as well as the relative taxonomy abundance. The weighted UniFrac beta diversity indices were calculated, and the microbial community structure differences were visualised with principal coordinate analysis plots (PCoA).

Data were visualised using R (version 4.5.2), ggplot2 (version 4.0.1), tidyverse (version 2.0.0), qiime2R (version 0.99.6) and ggprism (version 1.0.7).

#### Microbiome analysis of shotgun metagenomic sequences

Upon completion of sequencing, the Trim Galore pipeline (V0.6.10) [23] was employed for adapter trimming and quality filtering of the paired-end reads, with a quality threshold of 25 and a minimum read length of 50 bp. To remove the human genome from quality-controlled sequencing, hg38 was obtained from the UCSC Genome Browser [24] and mapped with Bowtie2 (version 2.5.4) [25]. Metagenomic assembly was carried out using MEGAHIT (v1.2.9) [26].

Taxonomic annotations were assigned to FASTQ reads using Kraken2 (version 2.17.1) against the PlusPF database, which contains Standard plus RefSeq protozoa & fungi [27]. Bracken (v.3.0.1) [28] was used to re-estimate bacterial abundances at taxonomic levels.

eggNOG-mapper (version 2.1.13) [29,30] was used to annotate the prokaryotic genome. The results were processed with KEGGaNOG (version 1.1.19) [31]. and the outputs were used by SHORTBRED (v. 0.9.3) [32] in conjunction with pre-computed shortbread markers to determine the presence and absence of virulence factors and antimicrobial resistance genes across the samples against the virulence factor database (VFDB) [33] and the Comprehensive Antibiotic Resistance Database (CARD) [33,34].

Metagenome-assembled genomes (MAGs) were co-assembled using all samples by MEGAHIT (v1.2.9) for binning. The co-assembled contigs were mapped with Bowtie2 (version 2.5.4) binned with MetaBAT2 [35]. MAGs were assigned to taxonomy by GTDB-Tk (version 2.6.1) [36–38].

## Results

### Microbial diversity and taxonomic profiles of soil at The Imperial Palace

After denoising and QC of reads derived from 16S rRNA amplicon sequencing by DADA2, approximately 80-100 thousand reads were obtained for each sample, as shown in **Supplementary Table 1**. After QC of reads derived from shotgun metagenomic sequencing by the Trim Galore pipeline, approximately 1.60–1.65 million base pairs were obtained for each sample (**Supplementary Table 2)**. To understand the taxonomic composition of soil at The Imperial Palace, our samples were classified at the phylum level by both 16S rRNA amplicon sequencing and metagenomic sequencing. The top eight phyla are shown in a stacked bar graph (**Figure 2A**) based on reads from shotgun metagenomic sequencing; the rest are noted as the remainder. Pseudomonadota (52.38 % ∼ 56.93 %) and Actinomycetota (17.39 % ∼ 34.89 %) were abundant in all samples. The top seven phyla are shown in a stacked bar graph (**Figure 2B**) based on reads from 16S rRNA amplicon sequencing; the remaining genera are noted as the remainder. At the phylum level, Proteobacteria, Acidobacteriota, and Actinobacteriota were abundant in almost all samples. In contrast to the other samples, KOKYO_01 exhibited a higher relative abundance of Bacteroidota. KOKYO_03 showed a relatively high abundance of Actinobacteriota and Chloroflexi. which are broadly consistent with the results of 16S rRNA amplicon sequencing. At the order level, Rhizobiales (1.61 ∼ 13.28 %), Burkholderiales (1.96 ∼ 18.24 %), Vicinamibacterales (0.55 ∼ 8.66 %), Gemmatimonadales (0.90 ∼ 7.64 %), and Acidobacteriales (0.01 ∼ 12.63 %) showed relatively higher abundance as shown in **Figure 2C**.

**Figure 2.**
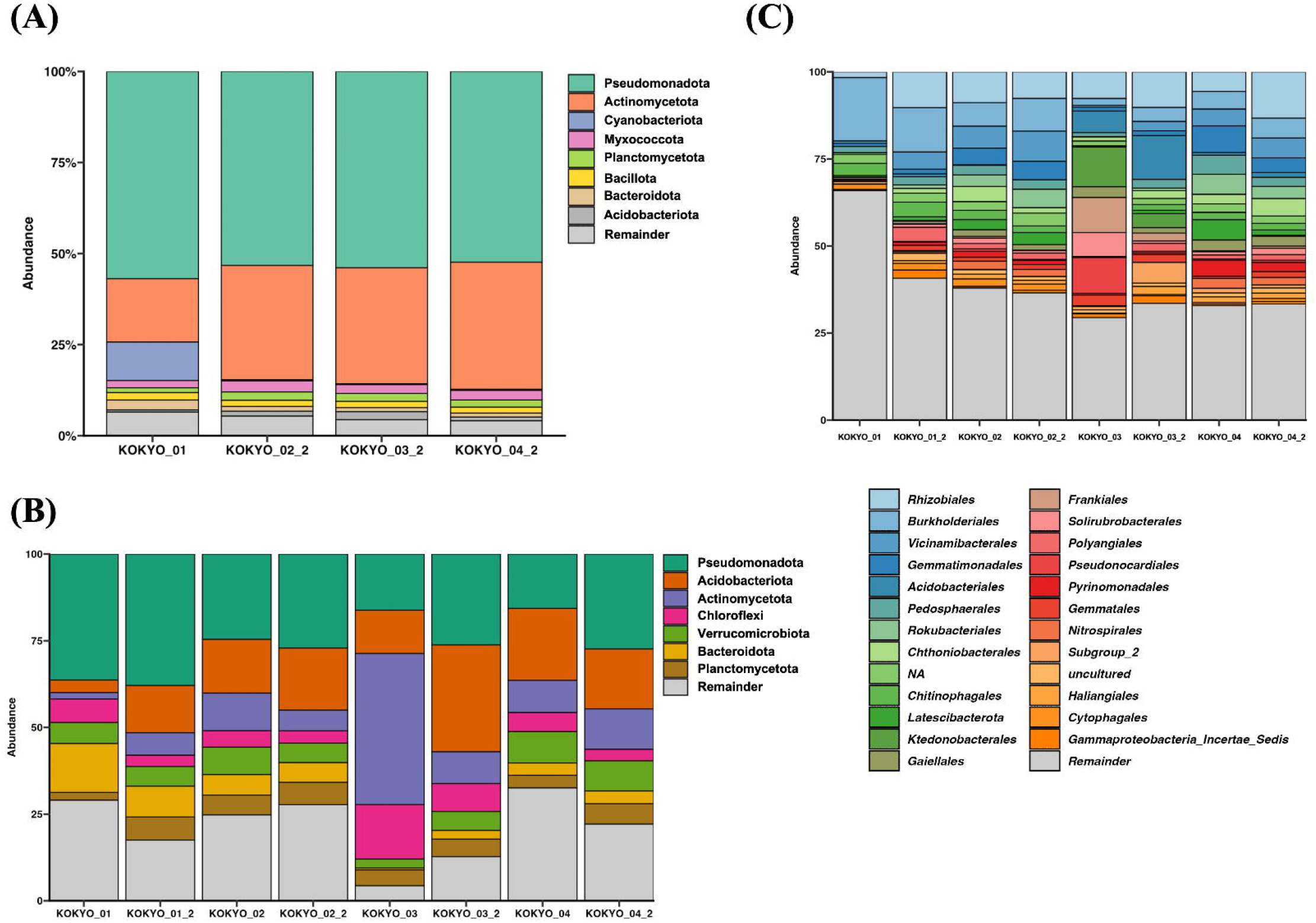
Relative abundance of most abundant microbial (A) phylum level based on the reads derived from metagenomic sequencing and (B) phylum level based on the reads derived from 16S rRNA amplicon sequencing. The ‘remainder’ represents the cumulative relative abundance of all other taxon not listed in the top 8 for each sample. (C) order level based on the reads derived from 16S rRNA amplicon sequencing. The ‘remainder’ represents the cumulative relative abundance of all other taxon not listed in the top 25 for each sample.

Furthermore, microbial diversity was evaluated for each sample. Principal coordinates analysis (PCoA) based on weighted UniFrac distances was performed (**Figure 3A**). KOKYO_02 and KOKYO_04 tended to have similar microbial structures. **Figure 3B and 3C** show the alpha diversity of each sample, assessed using Shannon index and Faith’s PD. In both metrics, KOKYO_03 samples (Shannon index, mean: 9.54; Faith’s PD, mean: 87.55) exhibited lower values compared to the other samples (Shannon index, mean: 10.18; Faith’s PD, mean: 125.55).

**Figure 3.**
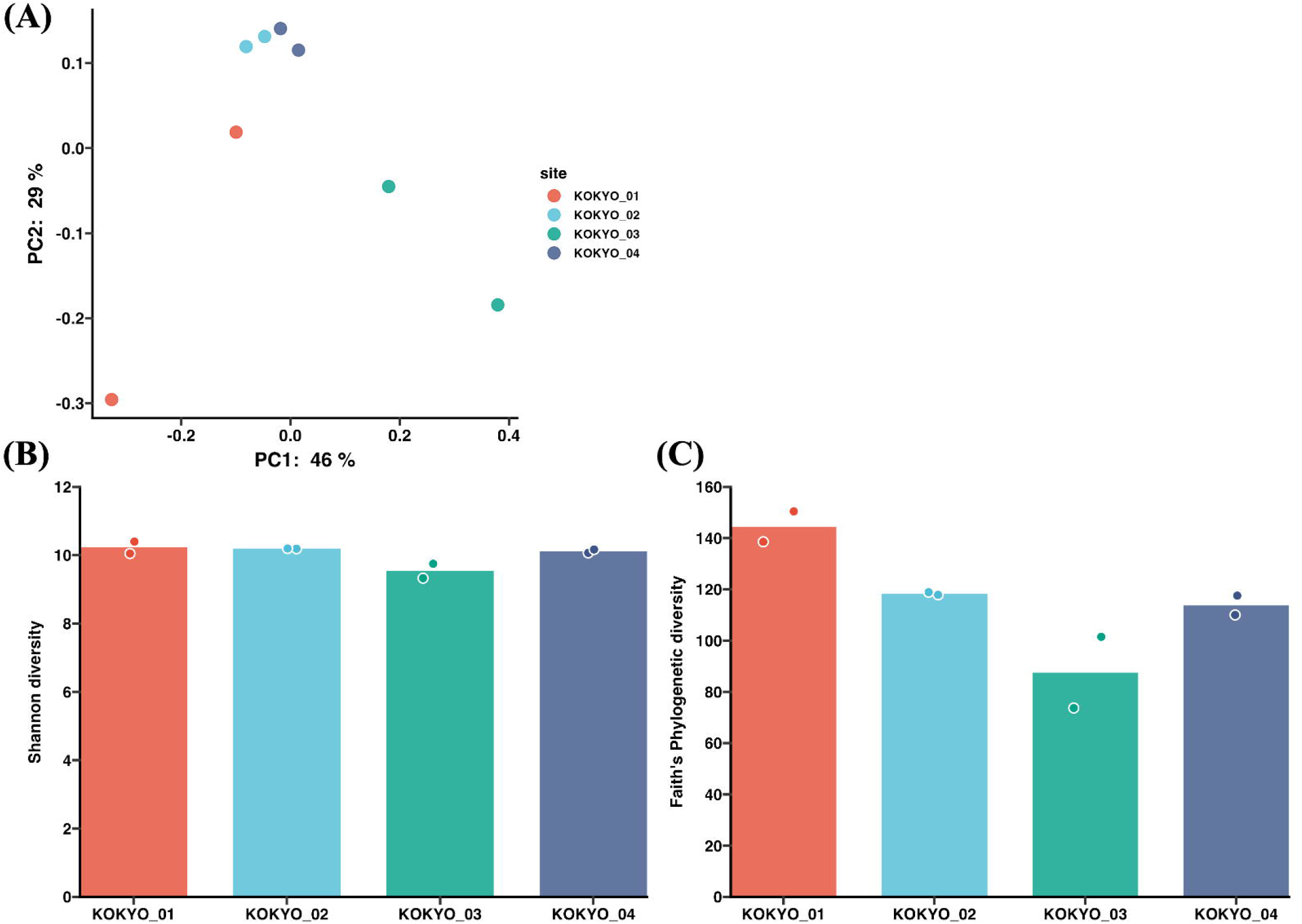
(A) Weighted Unifrac Principal Coordinate Analysis (PCoA) of samples analysed by 16S rRNA amplicon sequencing. (B) The Shannon diversity index and (C) Faith’s PD index for all samples based on the reads derived from 16S rRNA amplicon sequencing.

To identify uncultured and previously unknown microbial species, four shotgun metagenomic samples were co-assembled, and the generated contigs were binned using MetaBAT2 to reconstruct MAGs. As a result, 13 MAGs were obtained. However, none of the MAGs met the criteria for high quality (completeness ≥ 90% and contamination < 5%) or medium quality (completeness ≥ 50% and contamination < 10%) (**Supplementary Table 3**). The quality of the reconstructed MAGs is also summarized in **Supplementary Table 4**. Five MAGs (bin4, bin8, bin9, bin10, bin12) could not be classified at the domain level.

Several other MAGs were assigned only to higher taxonomic ranks, such as order or family, but could not be resolved at the genus or species level: bin5 to the family of Burkholderiaceae, bin6 to the family of HGW-15, bin11 to the order of Gloeomargaritales, and bin13 to the family of Competibacteraceae. The remaining bins were assigned to genus or species level: bin1 to the genus of *20CM-2-65-7*, bin2 to the genus of *Cyanobium*, bin3 to the genus of *QIAW01*, bin6 to the species of *Aquariibacter lacus*.As shown in **Supplementary Table 4**, the recovered bins accounted for an average of 0.68% across all samples, while over 99% of the genomes remained unassigned to these bins.

### Functional diversity of soil at The Imperial Palace

Based on the microbial community diversity observed in the Imperial Palace samples, we next sought to understand the resulting diversity of genetic functions. To obtain more robust gene annotations, co-assembly was performed across all samples. Gene annotation using eggNOG-mapper identified 248,570 genes, of which 81 were annotated as having unknown functions.

The completeness of metabolic pathways inferred from the annotated genes is illustrated in the heatmap shown in **Table 1**. Pathways exhibiting high completeness included “amino acid metabolism” (0.97), “carbohydrate metabolism” (0.96), “carbon fixation” (0.95), and “vitamin biosynthesis” (0.95). In contrast, pathways related to “biofilm formation” (0.22) and “genetic competence” (0.33) showed low completeness.

**Table 1.**
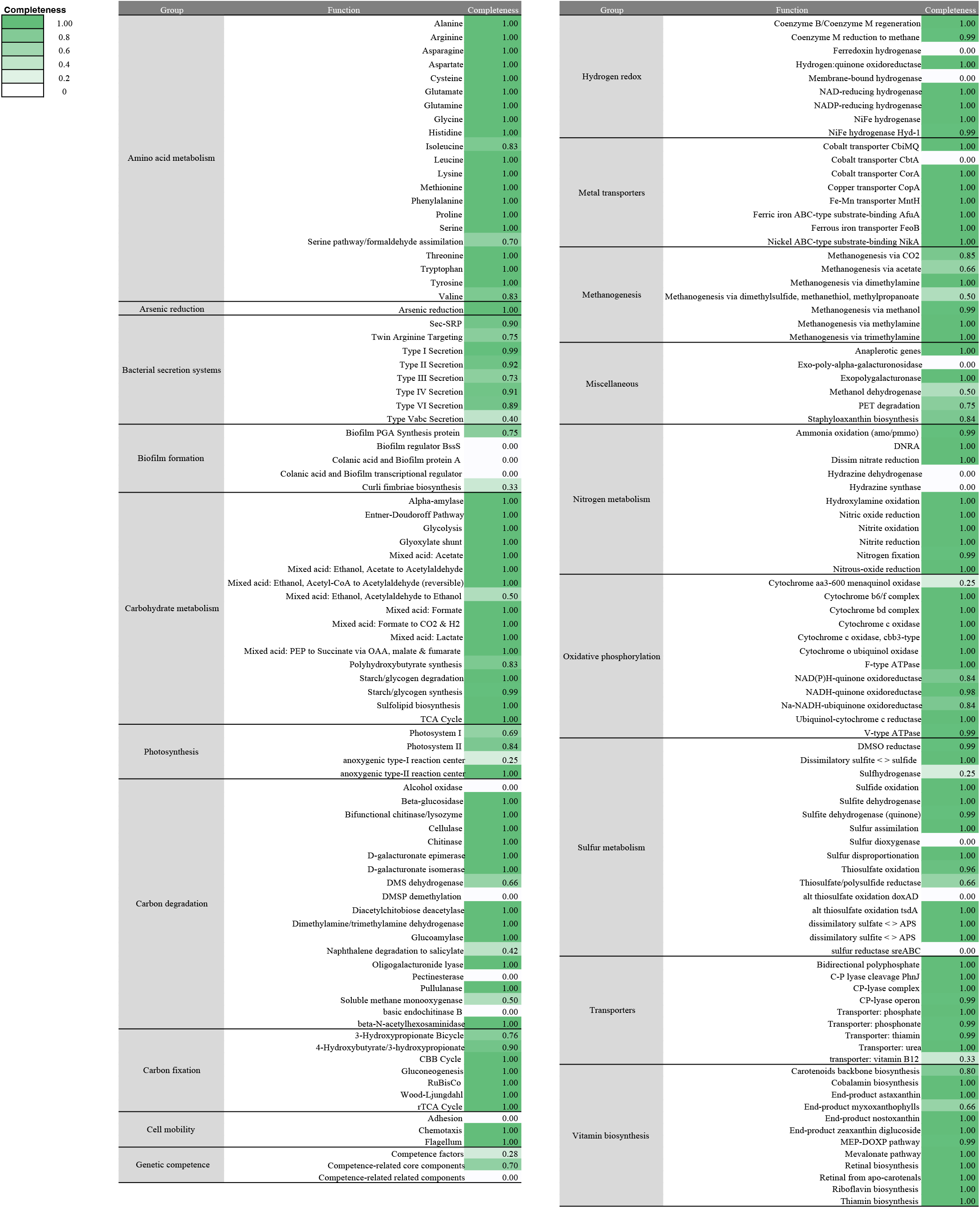
Heatmap illustrating the presence or absence of genes involved in various metabolic functions.

Next, the presence of genes associated with antimicrobial resistance and virulence was assessed using ShortBRED based on VFDB and CARD. As shown in **Supplementary Table 5**, no hits were detected in KOKYO_01, KOKYO_02_2, or KOKYO_04_2 against both VFDB and CARD. In contrast, KOKYO_03_2 contained hits to genes annotated as aminoglycoside resistance genes (virulence|440|vfid|441|vsiid|440|ssid|orf5_aadB) as well as genes encoding transposases (virulence|961|vfid|962|vsiid|961|ssid|transposase, virulence|961|vfid|68779|vsiid|91540|ssid|transposase), which were annotated within the virulence database.

## Discussion

To the best of our knowledge, this study represents the first genome-based analysis of microbial community structure in soils from the Imperial Palace in Japan. The microbial taxa that were abundant in the soil samples included members of the Rhizobiales [39,40], Burkholderiales [40–42], Vicinamibacterales [42], Gemmatimonadales [43], and Acidobacteriales [39], which are known as microbial taxa commonly found in soil and play vital roles in plant growth. For example, Rhizobiales is associated with nitrogen fixation, plant pathogenicity, and organic matter decomposition [44]. Burkholderiales is reported to suppress a broad spectrum of plant diseases caused by bacteria, fungi, oomycetes, and viruses while promoting plant growth [45]. Vicinamibacterales is a potential taxon contributing to the availability of soil phosphorus stock [46].

Focusing on microbial diversity, KOKYO_02 and KOKYO_04 tended to have similar microbial structures. This may be due to the soil of these samples originating from similar environments (mountainous soil with multiple deciduous trees). KOKYO_03 and KOKYO_03_2 exhibited relatively lower alpha diversity indexes (**Figure 3B, 3C**). The environment surrounding them was a managed garden composed primarily of a lawn dominated by a single plant species (**Figure 1C**), with Japanese black pine as the only tree species. This resulted in lower plant and tree diversity compared with the other sampling sites, which were characterized by diverse undergrowth and leaf litter accumulation. Although this observation is based on a limited number of samples, previous studies have reported that soil microorganisms metabolize soil nutrients into forms accessible to plants and that reciprocal interactions exist between the soil microbiome and plant growth [51–54]. Therefore, the monoculture-dominated vegetation at the site may represent one factor associated with the reduced microbial alpha diversity observed, as reported in the previous research [47–50].

Furthermore, the fact that samples with shared characteristics—such as diverse undergrowth and leaf litter accumulation (e.g., KOKYO_02 and KOKYO_04)—showed similarities in taxonomic profiles in the PCoA plot **(Figure 3A)** may further imply a functional interaction between plants and soil microbiome. Further studies incorporating a larger number of samples and explicitly linking microbial diversity to plant species richness are warranted to validate this relationship.

It should be noted that more than 98% of the genomic sequences in all samples were not mapped to the reconstructed bins. Although insufficient sequencing depth should be considered as a possible contributing factor, given the well-known tendency for MAGs to represent the most abundant taxa within a sample, the low number of mapped MAGs likely reflects the presence of highly diverse soil bacterial communities at the sampling sites, resulting in the formation of complex microbial communities [55].

As shown in **Table 1**, Functional gene analysis revealed the frequent presence of core metabolic pathways, including those involved in amino acid and carbohydrate metabolism across all samples. In contrast, pathways related to biofilm formation and genetic competence were rarely detected.

A limitation of this study is insufficient sequencing depth, which may prevent the recovery of high-quality MAGs. A further understanding of the complex soil microbiome requires greater sequencing depth.

## Conclusion

Thus, metagenomic analysis revealed, for the first time, the soil microbial community structure of the Imperial Palace Outer Gardens, demonstrating that Pseudomonadota and Actinomycetota were the predominant phyla. The soil samples collected from sites with different surrounding vegetation exhibited different levels of microbial diversity. Furthermore, only a small number of MAGs were mapped, which may reflect the presence of diverse soil bacterial communities at the sampling sites, resulting in the formation of complex microbial communities. Understanding the previously overlooked microbial diversity within the Imperial Palace Outer Gardens, a unique urban space where ecosystems remain highly preserved despite its central metropolitan location, can provide a valuable foundation for developing urban ecological frameworks and enhancing biodiversity conservation strategies.

## Acknowledgements

We would like to express our sincere gratitude to the Kokyogaien National Garden Management Office, Ministry of the Environment, for granting us permission to conduct sampling, and to the members of Nikken Sekkei Ltd. for their support of this research. This study was conducted as part of a research project aimed at enhancing the health and well-being of workers by incorporating natural elements into the Palaceside Building, a historic structure located near the Imperial Palace.

## Conflicts of interest

The authors are affiliated with BIOTA Inc. and declare no conflicts of interest relevant to this work.

## Author contributions

H.A. and K.I. designed the study and collected the samples. K.I. analysed the data. H.A. and R.F. interpreted the data. H.A was the major contributor in writing the manuscript. All authors read and approved the final manuscript.

## Figures and tables

**Supplementary Table 1**. Summary of 16S rRNA amplicon sequencing statistics.

**Supplementary Table 2**. Summary of metagenome sequencing statistics.

**Supplementary Table 3**. The quality of each bacterial MAGs reconstructed from all samples combined.

**Supplementary Table 4**. Relative abundance of MAGs for each sample.

**Supplementary Table 5**. Virulence factors and antibiotic resistance genes identified in the samples. ‘Family’ represents the database family and gene identifier, ‘count’ is the total abundance count of each gene, ‘hits’ is the number of times the gene was detected, and ‘marker length’ is the length of the gene marker.

